# Protracted abstinence from heroin, but not cocaine, is associated with profound medial prefrontal cortex synaptic bioenergetic remodeling and lasting neural hypofunction

**DOI:** 10.64898/2026.08.18.743579

**Authors:** Yun Young Yim, Romain Durand de Cuttoli, Tamara Markovic, Angelica Minier-Toribio, Arthur Godino, Freddyson J. Martínez-Rivera, Rita Futamura, Joseph Landry, Annie Ly, James E. Callens, Scott J. Russo, Yasmin L. Hurd, Angus C. Nairn, Eric J Nestler, Caleb J Browne

## Abstract

Relapse following prolonged abstinence is a primary challenge in the treatment of opioid and cocaine use disorders, driven in part by enduring dysfunction of medial prefrontal cortex (mPFC) circuits that impair inhibitory control over drug-seeking. The molecular substrates underlying this dysfunction, and whether they differ across drug classes, remain unknown. Here, we performed label-free quantitative proteomics of mPFC synaptosomes isolated from rats after 30-day abstinence following intravenous heroin or cocaine self-administration to profile synaptic adaptations that may contribute to relapse vulnerability. Heroin abstinence induced extensive synaptic proteomic remodeling characterized by coordinated downregulation of mitochondrial proteins involved in oxidative phosphorylation, including pyruvate dehydrogenase complex subunits that regulate carbon entry into mitochondrial metabolism. Targeted metabolomic profiling of whole mPFC revealed accumulation of upstream glycolytic and pentose phosphate pathway intermediates, consistent with altered pyruvate utilization and mitochondrial oxidation. Several bioenergetic metabolites also correlated positively with the severity of escalation of heroin intake. Consistent with the bioenergetic remodeling observed during protracted heroin abstinence, whole-cell patch-clamp recordings from layer V mPFC pyramidal neurons revealed lasting suppression of intrinsic excitability and a decreased spontaneous excitatory synaptic activity. Cocaine abstinence, by contrast, produced limited changes in synaptic bioenergetics while inducing a distinct cytoskeletal remodeling signature. Overall, these findings identify synaptic bioenergetic remodeling as a previously underappreciated feature of prolonged heroin abstinence and reveal a marked divergence in the molecular adaptations induced by heroin versus cocaine within the mPFC. These results implicate mitochondrial bioenergetic pathways as therapeutic targets for reducing relapse vulnerability specifically associated with opioid use disorder.

## Introduction

Substance use disorders exact a devastating toll on individuals, their families, and the healthcare system as a whole. Among them, opioid use disorder (OUD) and cocaine use disorder (CUD) account for a substantial proportion of drug overdose deaths and remain major causes of morbidity and mortality^1^. Although many individuals seek treatment and achieve periods of abstinence, relapses remain the defining clinical challenge, occurring in the majority of patients across treatment modalities^2–4^. Sustained abstinence without pharmacological intervention is uncommon in OUD^5^, and despite the effectiveness of opioid replacement therapies such as methadone and buprenorphine in reducing opioid use and overdose risk, relapse frequently occurs following treatment discontinuation^6–10^. In contrast, no pharmacological treatments are approved in the U.S. for CUD, leaving behavioral interventions as the sole therapeutic option despite their limited long-term efficacy^11^. This clinical intractability points to underlying neurobiological vulnerabilities that are not adequately addressed by current therapeutic strategies. Identifying the neurobiological mechanisms that maintain relapse vulnerability after prolonged abstinence therefore remains a critical challenge for the development of more effective interventions.

Relapse vulnerability is thought to arise from prolonged neuroadaptations within the brain reward circuitry that persist well beyond the period of active substance use^12–14^. Among the brain regions comprising the reward circuitry, the medial prefrontal cortex (mPFC) is well-positioned to exert top-down control over mesolimbic dopamine signaling and support executive functions that promote adaptive behavior including decision-making, inhibitory control, behavioral flexibility, and motivation^15–17^. Consistent with this role, neuroimaging and neuropsychological studies in individuals with OUD and CUD have demonstrated persistent prefrontal hypofunction during abstinence, characterized by reduced activation of the mPFC and anterior cingulate cortex during cognitive tasks, together with lasting deficits in attention, working memory, and inhibitory control^15, 18–21^. These impairments are thought to weaken executive control over substance-seeking behavior, thereby increasing vulnerability to relapse. Converging evidence from rodent models further demonstrates that selective manipulation of mPFC activity or its projections to the mesolimbic dopamine system causally regulates opioid- and cocaine-seeking behaviors with both activation and inhibition of specific mPFC subregions and their efferent pathways bidirectionally modulating drug seeking and extinction^22–27^. These findings converge on a model in which persistent mPFC dysfunction during abstinence interferes with top-down inhibitory control over substance-seeking circuits, thereby promoting relapse vulnerability.

The sustained suppression of mPFC function during prolonged abstinence may ultimately be encoded at the molecular level through durable adaptations in the protein machinery and structural organization of synapses^28, 29^. Long-lasting changes in neural circuit function can involve persistent remodeling of the synapses, including alterations in receptor subunit composition, scaffolding protein abundance, and vesicle release machinery^30–32^. Importantly, because synaptic transmission is among the most energetically demanding processes in the brain^33, 34^, maintaining cortical activity requires not only appropriate synaptic protein composition but also tight coordination with cellular metabolism and mitochondrial energy production^35^. Defining these enduring molecular adaptations within the mPFC synaptic compartment is therefore essential for understanding how relapse-associated brain states are established and maintained long after substance exposure has ceased.

Previous transcriptomic and proteomic studies have begun to characterize substance-induced neuroadaptations within reward-related brain regions, including transcriptome-wide analyses of heroin and cocaine self-administration^36, 37^ and targeted proteomic studies of the mPFC following heroin self-administration^38–40^. While these studies have provided important mechanistic insight, several critical gaps remain. Few studies have comprehensively examined the synaptic proteome after 30 days of abstinence, leaving the molecular machinery supporting lasting mPFC dysfunction largely undefined. Moreover, it remains unknown whether prolonged abstinence from opioids or psychostimulants engages shared or distinct molecular adaptations within the mPFC^41^, an important question for determining whether relapse vulnerability is maintained by convergent or drug-specific biological mechanisms.

To address these questions, in this study, we combined unbiased quantitative proteomic profiling of isolated mPFC synaptosomes with targeted metabolomic analysis and *ex vivo* electrophysiological recordings in rats following 30 days of forced abstinence from heroin or cocaine self-administration. This time point was selected to correspond to the well-established incubation of drug craving, during which substance seeking progressively increases and relapse vulnerability is heightened^42–45^. By integrating changes in the synaptic proteome with alterations in cellular metabolism and neuronal physiology, we sought to determine whether prolonged heroin and cocaine abstinence engage shared or distinct molecular mechanisms underlying persistent mPFC dysfunction. We further explored the biological pathways linking synaptic molecular remodeling to altered neuronal function and determined whether these adaptations reveal common mechanisms of relapse vulnerability or drug-specific molecular signatures that may inform future therapeutic strategies.

## Materials and methods

### Animals

Adult male Long-Evans rats (∼300 g) were maintained on a 12 h reverse light-dark cycle with food and water *ad libitum*, except during intravenous drug self-administration (IVSA) procedures (see below). All experiments were conducted during the dark phase. All animal procedures were conducted in accordance with the NIH Guide for the Care and Use of Laboratory Animals and approved by the Institutional Animal Care and Use Committee at the Icahn School of Medicine at Mount Sinai.

### Drugs

Heroin (diacetylmorphine HCl; contribution from NIDA) and cocaine (cocaine HCl; contribution from NIDA) were dissolved in 0.9% sterile saline.

### Intravenous drug self-administration

Rats received jugular vein catheterization under inhaled isoflurane anesthesia (2%). Rats were pair-housed prior to surgery and single-housed following surgery for the duration of the experiment. Following 1 week of recovery, rats were food restricted (18 g/day) and trained to self-administer heroin (0.03 mg/kg/inf) or cocaine (0.8 mg/kg/inf) via indwelling catheters, or saline as a control. Rats were tested in operant chambers (Med Associates Inc, St Albans, VT) housed within sound-attenuating cabinets. Chambers were illuminated by a house light and two response levers were extended. Responding on the active lever resulted in a 5 s infusion of drug or saline concomitant with retraction of both levers, extinguishing of the house light, and illumination of two cue lights for 20 s according to a fixed-ratio 1 schedule of reinforcement. After the 20 s timeout, the house light was re-illuminated, cue lights were extinguished, and levers were re-inserted into the chamber. Responding on the inactive lever had no programmed consequence. Drug delivery was controlled by a syringe pump located outside of the sound attenuating cubicles. Each self-administration session lasted 6 h and rats underwent 10-15 sessions based on patterns of acquisition defined as a minimum of 20 infusions per session of either heroin or cocaine earned; Session 1 was considered the first session in which rats met this criterion. Following the final IVSA session, rats were returned to *ad libitum* food access and remained in home cages undisturbed for 30-day forced abstinence. On day 30, rats were euthanized by decapitation followed by rapid brain extraction and tissue dissection on ice. Brains were sectioned on a 1 mm thickness brain matrix, and mPFC punches were collected using a 12 g needle, spanning both prelimbic and infralimbic mPFC, and immediately flash frozen. Behavioral data were analyzed using two-way ANOVAs comparing either heroin or cocaine to saline, with group effects.

### Synaptosome preparation

Crude synaptosomes were prepared from bilateral mPFC punches using previously described protocols^46^. Briefly, mPFC tissue was homogenized in 400 μL of HEPES-sucrose buffer containing 0.32 M sucrose, 4 mM HEPES (pH 7.4), protease inhibitors (Roche cOmplete Mini), and phosphatase inhibitors (Roche PhosSTOP). Homogenates were centrifuged at 1,000 × g for 10 min at 4°C, and the resulting supernatants (S1 fraction) were collected. Pellets were resuspended in 400 μL of fresh HEPES-sucrose buffer and centrifuged again under the same conditions, and the supernatants were pooled. The combined S1 fraction was then centrifuged at 10,000 × g for 10 min at 4°C to obtain the crude synaptosomal pellet. This pellet was resuspended in 400 μL of fresh HEPES-sucrose buffer and centrifuged once more under the same conditions. After discarding the final supernatant, the resulting crude synaptosomal pellet (P2) was snap-frozen and stored at −80 °C for subsequent proteomics analysis.

### Synaptosome Proteomics

Proteomics analysis was conducted at the Yale/NIDA Neuroproteomics Center Core. Synaptosomes were dissolved in 8 M urea / 0.4 M ammonium bicarbonate (ABC), and 20 µL of each sample was processed for reduction and alkylation, followed by trypsin digestion and desalting. All samples were analyzed by label-free quantification in DIA mode, using targeted feature extraction based on PFC-specific spectral libraries generated from DDA runs of in-house rat synaptosome samples. To support targeted feature extraction, a PFC-specific spectral library was generated from DDA analyses of synaptosomal samples prepared from naïve Long-Evans rats maintained under comparable housing conditions. The resulting mass spectrometry proteomic data have been deposited in [repository name] under accession number [accession ID].

### Proteomic analysis

Proteomic data were processed using Scaffold DIA (v2.1.0; Proteome Software). Protein abundances were quantified from normalized exclusive peptide intensities using EncyclopeDIA within Scaffold DIA and exported as log_2_ fold-change values relative to the mPFC-saline group. Protein identification required a false discovery rate (FDR) of 1% and a minimum of two unique proteolytic peptides per protein. Only reviewed Swiss-Prot protein entries were retained for downstream analyses. Principal component analysis (PCA) was performed to assess sample clustering and identify potential outliers.

Differential protein abundance was analyzed in R (v4.3.1) using the limma package. Separate pairwise comparisons were performed for cocaine versus saline and heroin versus saline. For each comparison, linear models were fitted to protein abundance data, and variance estimates were moderated using the empirical Bayes procedure implemented in limma. The resulting statistics included log fold changes, moderated *t* statistics, nominal *P* values, Benjamini– Hochberg false discovery rate (FDR)-adjusted *P* values, and log-odds of differential abundance (B statistics). Proteins meeting the predefined significance criteria (absolute log_2_ fold change >0.263 and nominal limma p< 0.05) were considered differentially abundant and used for downstream pathway analyses. To assess the robustness of the observed group differences, permutation testing was additionally performed by randomly permuting sample group labels 1,000 times using a fixed random seed and repeating the limma analysis for each permutation. For each protein, an empirical *p* value was calculated as the proportion of permuted absolute moderated *t* statistics greater than or equal to the absolute observed moderated *t* statistic.

For visualization and generation of differential protein lists, proteins were classified as significantly altered if they exhibited an absolute log_2_ fold change >0.263 (corresponding to an approximately 20% change in abundance) together with a nominal limma p value < 0.05. Volcano plots were generated using the EnhancedVolcano R package, with log fold change plotted against the nominal p value. Proteins exhibiting an absolute log_2_ fold change >0.263 and a nominal p value <0.05 were classified as significantly, differentially abundant and displayed as increased or decreased, whereas all remaining proteins were classified as not significant. To identify representative proteins for annotation, proteins were ranked using a composite score calculated as the product of the absolute log fold change and the negative log of the nominal *p* value (|log_2_FC| × −log_10_[*p*]). The three increased and three decreased proteins with the highest composite scores were labeled in each volcano plot. Heatmaps were generated using Morpheus (Broad Institute). Gene Ontology enrichment analysis was performed using Enrichr using the 2025 Gene Ontology Biological Process, Molecular Function, and Cellular Component databases. Ingenuity Pathway Analysis (IPA; Qiagen) was used to identify enriched canonical pathways and predict upstream regulators. Protein protein interaction networks were generated using STRING^47^ (https://string-db.org/).

### Metabolomics

#### Sample processing

Metabolomics analysis was conducted at the NYU Metabolomics Core. Metabolites were extracted from mPFC punches of rats following 30-day forced abstinence from self-administered cocaine, heroin, or saline using a hybrid LC-MS assay as described previously^48^. Briefly, tissue samples were extracted in 80% methanol, 1% formic acid, and 500 nM metabolomics amino acid mix internal standard (catalog no. MSK-A2-1.2, Cambridge Isotope Laboratories) (5 mg tissue/mL buffer), followed by bead homogenization (10 cycles of 30 s at 6 m/s with 30 s pauses) and centrifugation (21,000 × g, 3 min, 4°C). Supernatants were dried by SpeedVac, reconstituted in LC/MS-grade water, sonicated for 2 min, and centrifuged (21,000 × g, 3 min, 4°C) prior to LC-MS analysis. Samples were analyzed by LC-MS/MS using a Millipore ZIC-pHILIC column (2.1 × 150 mm, 5 μm) maintained at 25°C, coupled to a Dionex Ultimate 3000 UHPLC and a Thermo Q Exactive HF mass spectrometer operating in HESI mode. Separation was performed using a gradient of 10 mM ammonium carbonate (pH 9.0) (buffer A) and acetonitrile (buffer B) at 100 μL/min over a 42-min run (80-20% B, 0-30 min; 20-80% B, 30-31 min; 80% B, 31-42 min; 2 μL injection volume). Data were acquired in positive and negative ionization modes using data-dependent MS/MS (Top 5) with polarity switching. Full-scan spectra (m/z 67-1000; 120,000 resolution) and tandem MS spectra (15,000 resolution; multiplexed normalized collision energies of 10, 35, 80) were collected in profile mode for metabolite identification and quantification.

#### Data Analysis

Raw LC-MS/MS data were converted to SQLite format and processed using in-house Python-based pipelines for peak detection and quantification (https://github.com/NYUMetabolomics/plz). Metabolites were identified by matching accurate mass, retention time, and MS/MS spectra against an internally curated library developed using authentic standards and validated against NIST14 and METLIN (2017) databases^49, 50^. Peak intensities were extracted using a ±15 ppm mass tolerance and a ±7.5 s peak apex retention time tolerance, within an initial retention time search window of ±0.5 min and normalized using a cocktail of 13C/15N-labeled amino acid internal standards. Instrument performance was monitored through mass accuracy, retention time stability, and internal standard reproducibility, with median mass error ≤10 ppm, retention time variation ≤0.5 min, and internal standard CVs generally ≤15%. Peak intensities were blank-corrected using an in-house statistical pipeline (Metabolyze v.1.0) and filtered based on a signal-to-noise ratio ≥3, with a floor of 10,000 (arbitrary units); metabolites below this threshold were annotated as not detected, and the threshold value was imputed for statistical comparisons to enable estimation of fold change where applicable. Based on findings from synaptic proteomics showing alterations in cellular energetic pathways, statistical comparisons focused a priori on detectable metabolites within glycolysis, the pentose phosphate pathway, and the TCA cycle. T-tests were performed using SciPy (Python, 1.1.0), with metabolites exhibiting p< 0.05 considered significantly regulated.

#### Correlation with escalation of intake

To confirm that escalation of drug intake occurred at the group level, heroin and cocaine intake across sessions 3–10 (excluding sessions 1-2 to minimize acquisition effects) were each assessed using a linear mixed-effects model with session as a fixed effect and subject-specific random intercepts and slopes. Individual escalation slope was then derived for each animal to relate escalation magnitude to metabolite abundance. Escalation was quantified per animal by fitting a simple linear regression model with intake as the dependent variable and session as the independent variable; the resulting per-session slope was multiplied by the 7-session span to estimate each animal’s total change in intake across the escalation window. Metabolites included in the correlation analysis were restricted to those reliably detected in all samples, and intensities were z-scored row-wise across animals (mean = 0, SD = 1). Within each drug group (heroin, cocaine) separately, Spearman rank correlations (ρ) were computed between each animal’s escalation slope and each metabolite’s z-scored abundance (two-sided, asymptotic approximation). Multiple comparisons were controlled within each drug group using the Benjamini-Hochberg false discovery rate procedure (FDR, α = 0.05). All analyses were performed using R.

### Electrophysiology

Rats were decapitated, brains were rapidly extracted on ice, and coronal slices (250 µm thick) were prepared using a Compresstome (VF-300; Precisionary Instruments) in cold (0–4°C) sucrose-based artificial cerebrospinal fluid (aCSF) containing: 87 mM NaCl, 2.5 mM KCl, 1.25 mM NaH2PO4, 4 mM MgCl2, 23 mM NaHCO3, 75 mM sucrose, and 25 mM glucose. After 15 min incubation at 35°C for recovery, slices were transferred into oxygenated (95% O2/5% CO2) aCSF containing: 130 mM NaCl, 2.5 mM KCl, 1.2 mM NaH2PO4, 2.4 mM CaCl2, 1.2 mM MgCl2, 23 mM NaHCO3, and 11 mM glucose at room temperature, then transferred to a recording chamber continuously perfused at 2-3 mL/min with oxygenated aCSF. Patch pipettes (4-7 MΩ) were pulled from thin wall borosilicate glass using a micropipette puller (P-97, Sutter Instruments) and filled with a K-Gluconate based intra-pipette solution containing: 116 mM K-Gluconate, 20 mM HEPES, 0.5 mM EGTA, 6 mM KCl, 2 mM NaCl, 4 mM ATP, 0.3 mM GTP and 2 mg/mL biocytin (pH adjusted to 7.2). Cells were visualized using an upright microscope with an IR-DIC lens and illuminated with a white light source (Scientifica). Whole-cell recordings of Layer V mPFC pyramidal neurons were performed using a patch-clamp amplifier (Axoclamp 200B, Molecular Devices) connected to a Digidata 1550 LowNoise acquisition system (Molecular Devices). Excitability was measured in current-clamp mode by injecting incremental steps of current (0-300 pA, +20 pA at each step). For recording of spontaneous Excitatory Post-Synaptic Currents (sEPSCs), neurons were recorded in voltage-clamp mode at - 70 mV and events were detected with a 10 pA negative threshold. Signals were low-pass filtered (Bessel, 2 kHz) and collected at 10 kHz using the data acquisition software pClamp 11 (Molecular Devices). Electrophysiological recordings were extracted and analyzed using Clampfit (Molecular Devices). Neuronal excitability was analyzed using a two-way repeated-measures ANOVA with injected current (0-300 pA) as the repeated-measures factor and treatment as the between-group factor, followed by Šídák’s multiple-comparison test for comparisons between groups at individual current steps. Spontaneous excitatory postsynaptic current (sEPSC) frequency was compared between heroin- and saline-abstinent animals using Welch’s *t*-test. Statistical analyses were performed using GraphPad Prism 10 (GraphPad Software), and statistical significance was defined as *p< 0.05 and **p<0.01.

## Results

To assess how molecular adaptations in mPFC synapses may contribute to dysfunction during protracted abstinence, we combined IVSA of heroin or cocaine with an unbiased proteomic screen of isolated synaptosomes (**Fig. 1A**). Rats underwent a long-access (6 h) IVSA of heroin (0.03 mg/kg/infusion) or cocaine (0.8 mg/kg/infusion) or saline on a fixed-ratio 1 schedule of reinforcement. Rats in heroin and cocaine groups demonstrated significantly higher intake and selective active lever responding compared to saline controls (**Fig. 1B**). Following IVSA, rats underwent 30 days of home-cage forced abstinence, at which point they were euthanized, mPFC tissue was collected, and synaptosomes (P2 fraction) were isolated. Representative transmission electron microscopy confirmed enrichment of the synaptosomal preparation (**Fig. 1C**). Subsequently, mPFC synaptosomes (n=7 for each group) underwent label-free quantification mass spectrometry to characterize proteomic changes associated with protracted abstinence from heroin or cocaine IVSA.

**Figure 1.**
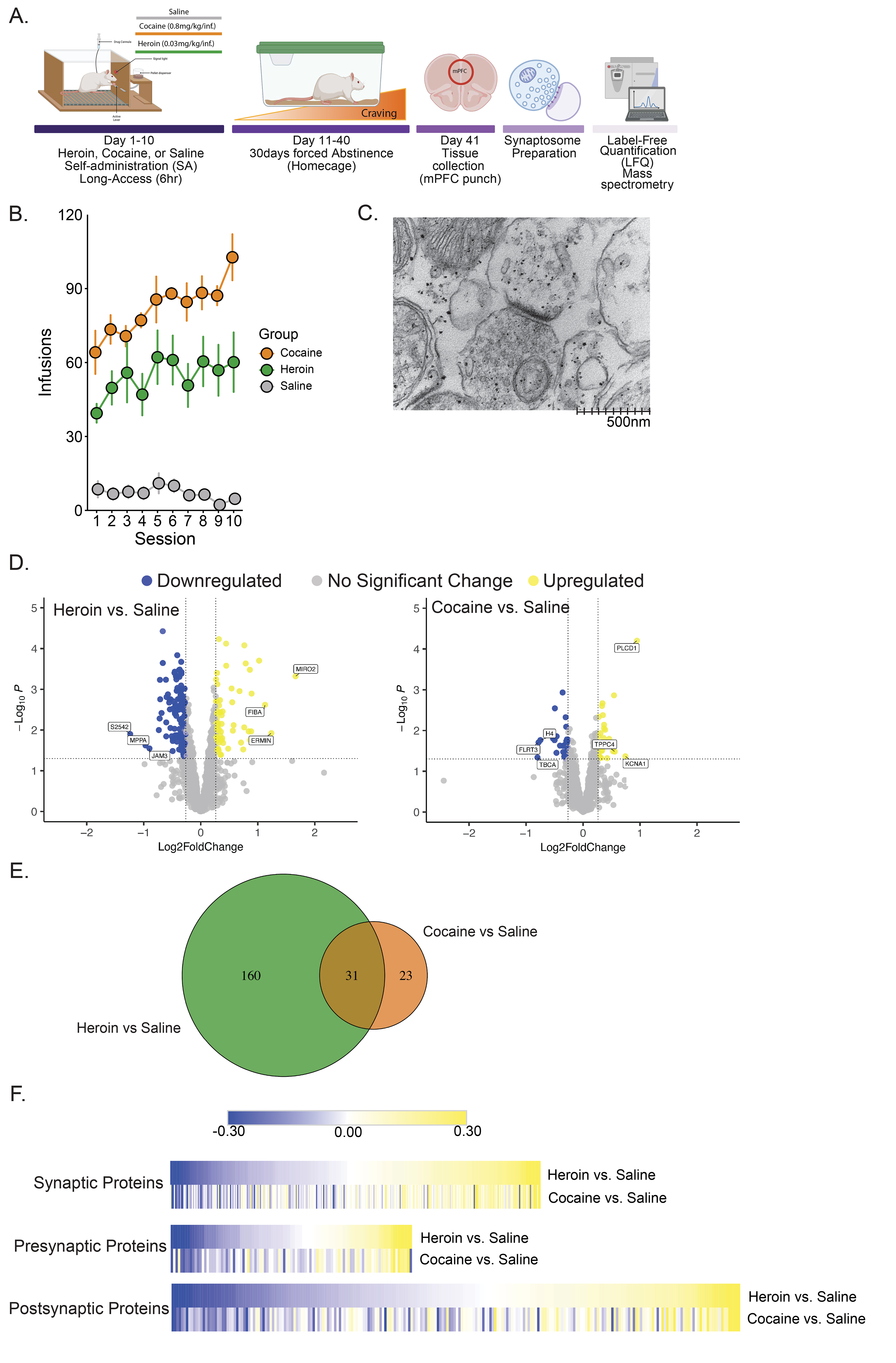
Long-access heroin and cocaine self-administration produce persistent remodeling of the mPFC synaptosomal proteome. **A.** Schematic outlining experimental design. Rats underwent 10 days of long-access intravenous self-administration (6 h/day) for heroin (0.03 mg/kg/infusion), cocaine (0.8 mg/kg/infusion), or saline, followed by 30 days of home-cage forced abstinence. Medial prefrontal cortex (mPFC) tissue was collected on abstinence day 30 for synaptosome isolation and label-free quantitative (LFQ) mass spectrometry analysis. **B.** Rats acquired self-administration behavior and showed significantly higher number of infusions for heroin (n=7; ANOVA, Group: F_(1,120)_=227.3, p< 0.001) or cocaine (n=7; ANOVA, Group: F_(1,120)_=1139.4, p< 0.001) compared to saline controls (n=7) (lower panel). **C.** Representative electron micrograph of an mPFC synaptosome preparation showing a presynaptic terminal adjacent to a postsynaptic density, demonstrating enrichment of synaptic structures in the crude synaptosomal preparation. **D.** Volcano plot showing protein abundance changes following heroin (left) or cocaine (right) abstinence relative to saline controls. Yellow indicates increased proteins and dark blue indicates decreased proteins, and grey indicates proteins that did not meet the differential-expression criteria of |log_2_FC|>0.263 and p< 0.05. **E.** Venn diagram showing number of differentially expressed proteins (DEPs) unique to and shared between the heroin vs. saline and cocaine vs. saline comparisons. **F.** Threshold-free heatmaps showing average log_2_ fold changes relative to saline for general synaptic, presynaptic, and postsynaptic protein categories defined using SynGo-annotations.

### Protracted heroin abstinence produces substantially greater synaptic proteomic remodeling in the mPFC compared to cocaine abstinence

Proteomic analysis revealed significant remodeling of the mPFC synaptic protein composition during protracted abstinence from heroin and cocaine, with heroin abstinence producing a substantially larger number of differentially expressed proteins (DEPs) than cocaine (191 vs. 54 DEPs; **Supplementary Table 1**). Principal component analysis (PCA) revealed treatment-associated differences in sample distribution, with heroin-abstinent samples showing greater differentiation from saline controls than cocaine abstinent samples, consistent with the broader proteomic remodeling observed following heroin abstinence. (**Supplementary Fig. 1A**).

Heroin abstinence was associated with a predominance of downregulated proteins (127 downregulated, 64 upregulated), whereas the cocaine proteome changes were more evenly distributed between downregulated and upregulated proteins (29 down, 25 up) (**Fig. 1D**). In addition, heroin-associated DEPs exhibited a broader range of fold changes than those identified following cocaine abstinence, indicating more extensive remodeling of synaptic proteome. Direct comparison of the two abstinence conditions identified 31 common DEPs (**Fig. 1E**). These overlapping proteins represented more than half of all cocaine-associated DEPs, indicating that many cocaine-associated changes occurred within a subset of the broader molecular adaptations observed following heroin abstinence. Nevertheless, the majority of heroin-associated DEPs were not shared with cocaine abstinence, highlighting substantial drug-specific remodeling of the mPFC synaptic proteome.

To compare the global synaptic proteomic landscape across drug classes, we generated threshold-free heatmaps illustrating all proteins annotated in the synaptic gene ontology (SynGO) database, organized into general synaptic, presynaptic, and postsynaptic protein categories (**Fig. 1F**). Across all three synaptic compartments, heroin and cocaine abstinence exhibited broadly similar directional changes for many synaptic proteins. However, heroin abstinence consistently produced larger average log_2_ fold changes across a greater number of proteins, indicating substantially more extensive synaptic proteomic remodeling than cocaine abstinence. Corresponding z-score heatmaps of normalized protein abundance across saline-, heroin-, and cocaine-abstinent groups (**Supplementary Fig. 1B**) further illustrate drug-dependent differences in synaptic protein abundance that underline the global patterns of proteomic remodeling observed in Fig. 1E.

### Synaptic proteomic changes reveal pervasive mitochondrial bioenergetic deficits during heroin abstinence

Gene ontology (GO) analysis of DEPs revealed distinct biological processes affected during protracted abstinence from heroin or cocaine intake. In heroin-abstinent rats (**Fig. 2A**, left; **Supplementary Table 2**), we observed a striking enrichment of downregulated proteins in GO terms, with the strongest engagement of biological processes including oxidative phosphorylation, ATP synthesis, and cellular respiration. These findings suggest heroin abstinence involves widespread suppression of proteins involved in mitochondrial bioenergetic pathways in the mPFC. In contrast, relatively few GO terms showed robust enrichment of upregulated proteins. The most significantly enriched biological process was glutathione-related processes, driven by increased expression of glutamate-cysteine ligase catalytic subunit (GCLC), the rate-limiting enzyme in glutathione biosynthesis^51^, together with the glutathione S-transferases μ3 (GSTM3) and Ω1 (GSTO1)(**Supplementary Table 2)**. These changes suggest engagement of glutathione-associated antioxidant pathways during heroin abstinence.

**Figure 2.**
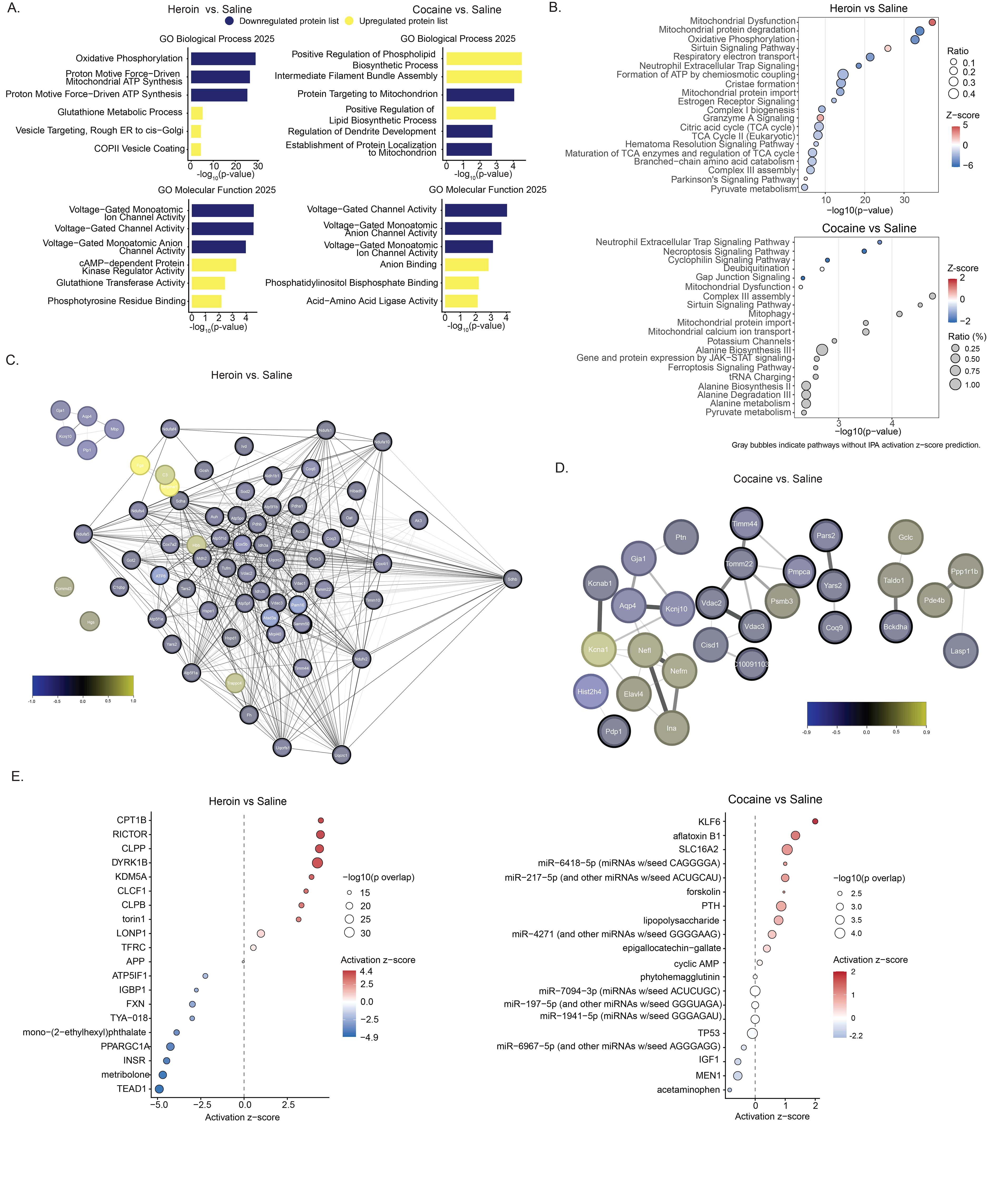
Heroin abstinence induces more extensive mitochondrial bioenergetic remodeling of the mPFC synaptosomal proteome than cocaine abstinence. **A.** Gene Ontology (GO) enrichment analysis of proteins differentially expressed following heroin or cocaine abstinence relative to saline control (|log_2_FC|>0.263, p<0.05) in the mPFC. The three most enriched biological process and molecular function categories among downregulated (blue) and upregulated (yellow) proteins are shown. Heroin abstinence was characterized by robust enrichment of mitochondrial respiration, ATP synthesis, and oxidative metabolic processes, whereas cocaine abstinence showed a more limited enrichment profile, including mitochondrial import among downregulated proteins and cytoskeletal/neurofilament-associated processes among upregulated proteins. **B.** Ingenuity Pathway Analysis (IPA) canonical pathway enrichment of DEPs. Bubble size indicates the ratio of proteins from the dataset mapped to each pathway, the x-axis represents pathway enrichment significance (-log10 p-value), and color indicates the predicted activation z-score. Heroin abstinence was associated with broad remodeling of pathways related to mitochondrial respiration, oxidative phosphorylation, TCA cycle, and cellular energy production. Cocaine abstinence was associated with fewer significantly enriched pathways and a larger proportion of pathways without a predicted activation state. **C-D.** String protein-protein interaction (PPI) network of DEPs following heroin (C) and cocaine (D) abstinence. Edges represent evidence-based protein associations at a minimum interaction score of 0.4. Node color indicates log_2_ fold change relative to saline, and proteins annotated with mitochondrial functions are highlighted by thicker node outlines. **C.** To reduce network complexity and emphasize the most robust changes, the heroin network was restricted to proteins meeting p< 0.01 with at least a 30%-fold change in abundance relative to saline controls (log_2_FC ≤ −0.515 or ≥ 0.379). The network reveals a highly interconnected cluster enriched for mitochondrial respiratory chain proteins, metabolic enzymes, and proteins involved in cellular energy production. **D.** The cocaine network included DEPs meeting |log_2_FC|>0.263 and p< 0.05 and exhibited smaller, more fragmented clusters involving mitochondrial protein import, astrocyte-associated proteins, ion-channel regulation, and neurofilament proteins. **E.** IPA upstream regulator analysis identifying candidate regulatory programs associated with the heroin (left)- and cocaine (right)-abstinent synaptosomal proteomes. Bubble size represents enrichment significance (-log10 overlap p-value), and color indicates the predicted activation z-score. Heroin abstinence was associated with predicted inhibition of regulators involved in mitochondrial biogenesis, respiratory function, and metabolic substrate utilization, including PGC-1α, FXN, and INSR, together with predicted activation of regulators associated with mitochondrial quality control and metabolic adaptation. Cocaine abstinence produced a more limited regulatory profile distinguished by predicted activation of KLF6 and identification of multiple candidate microRNA regulators. Collectively, GO enrichment, canonical-pathway analysis, STRING network organization, and upstream-regulator prediction converged on mitochondrial bioenergetic remodeling as the dominant molecular signature of 30-day heroin abstinence while suggesting distinct regulatory mechanisms following heroin and cocaine abstinence.

In cocaine-abstinent rats (**Fig. 2A**, right; **Supplementary Table 3**), GO analysis revealed a considerably more limited pattern of proteomic regulation than that observed during heroin abstinence. Downregulated proteins were enriched for mitochondrial protein import, driven by mitochondrial-processing peptidase subunit α (PMPCA), mitochondrial import inner membrane translocase subunit (TIMM44), and (TOMM22), key components of the mitochondrial translocase machinery^52^ (**Supplementary Table 3)**. While this finding indicates some overlap with the mitochondrial alterations observed following heroin abstinence, the broader suppression of bioenergetic pathways was not evident. Among upregulated proteins, intermediate filament bundle assembly was the most significantly enriched biological process, driven largely by the neurofilament subunits NFL and NFM. This enrichment is consistent with molecular remodeling of neurofilament and cytoskeletal pathways within mPFC axons. Additional support for this interpretation comes from increased expression of ELAV-like protein 4 (ELAVL4), an RNA-binding protein implicated in neuronal structural plasticity^53^, and the presynaptic scaffolding protein liprin-α3 (PPFIA3).

These GO analyses suggest changes to mitochondrial bioenergetic pathways as the defining feature of heroin abstinence while highlighting distinct upregulated signatures across drug classes, particularly in management of oxidative stress responses in heroin abstinence and cytoskeletal remodeling in cocaine abstinence. These distinct enrichment profiles suggest unique modes of synaptic adaptation in the mPFC during protracted abstinence from heroin and cocaine.

### Canonical pathway and protein interaction network analyses identify coordinated suppression of mitochondrial pathways during heroin abstinence

We next examined pathway-level alterations associated with synaptic proteomic remodeling using the Ingenuity Pathway Analysis (IPA) of DEPs (**Fig. 2B; Supplementary Table 4**). Consistent with GO enrichment analysis, IPA canonical pathway analysis confirmed extensive changes to mitochondrial pathways governing oxidative phosphorylation, respiratory chain function, oxidative metabolism, and mitochondrial assembly. The mitochondrial dysfunction pathway showed the strongest enrichment and a positive predicted activation z-score, whereas oxidative phosphorylation was predicted to be inhibited, providing convergent evidence for coordinated remodeling of mitochondrial bioenergetic pathways of mPFC synapses during heroin abstinence. Additionally, mitochondrial protein degradation, mitochondrial assembly, oxidative metabolism, and respiratory chain-related pathways were predicted to be inhibited, indicating broad disruption of mitochondrial function. In addition to these bioenergetic pathways, sirtuin signaling was also found to be significantly enriched and moderately activated. Sirtuins function as NAD+ dependent metabolic sensors that regulate mitochondrial function, oxidative metabolism, and cellular stress responses through reversible protein deacetylation^54^. Enrichment of this pathway therefore indicates remodeling of NAD+-dependent metabolic and stress response signaling concurrent with mitochondrial bioenergetic pathways during protracted heroin abstinence.

In contrast, cocaine abstinence exhibited considerably fewer enriched canonical pathways and smaller predicted activation states than heroin abstinence. Nevertheless, several mitochondrial-related pathways, including mitochondrial dysfunction, complex III assembly, mitochondrial protein import, mitophagy, and sirtuin signaling, overlapped with those identified in heroin-abstinent animals, suggesting limited engagement of mitochondrial regulatory pathways without the broad bioenergetic remodeling observed in heroin abstinent animals. Gap junction signaling was significantly inhibited and was consistent with downregulation of the gap junction protein connexin-43 (GJA1) at the protein level.

We then used STRING protein-protein interaction (PPI) analysis to further reveal how proteins altered during protracted abstinence were connected within functional interaction networks in mPFC synaptosomes (**Fig. 2C and D**). Consistent with GO and IPA analysis, heroin abstinent rats exhibited a larger, highly interconnected network of DEPs dominated by downregulated mitochondrial proteins (nodes with thick black outline) which formed the network’s core. This core network included proteins involved in respiratory chain function, mitochondrial trafficking, and energy metabolism, indicating coordinated suppression of bioenergetic machinery (**Fig. 2C**). Notably, this network included multiple subunits of the pyruvate dehydrogenase complex (PDHA1, PDHB)^55^ together with malate dehydrogenase 2 (MDH2)^56^, a key enzyme linking glycolysis to mitochondrial oxidative metabolism and regulating TCA cycle flux. In contrast, proteins altered following cocaine abstinence formed a smaller and more fragmented network (**Fig. 2D**). A discrete cluster of downregulated proteins associated with mitochondrial import was identified, consistent with the enrichment of mitochondrial pathways observed by GO and IPA analyses. These findings demonstrate that the coordinated bioenergetic pathways identified by GO and IPA are organized into highly interconnected mitochondrial protein networks within the synaptic proteome.

Interestingly, both heroin- and cocaine-abstinent animals exhibited downregulation of a network containing the canonical astrocytic proteins, aquaporin-4 (AQP4), inwardly rectifying potassium channel Kir4.1(KCNJ10), and connexin-43 (Gja1; **Fig. 2C and D**). In heroin-abstinent animals, this network appeared largely independent of the extensive mitochondrial network. In contrast, in cocaine-abstinent animals, these astrocyte-associated proteins were integrated within a network containing upregulated neurofilament protein, consistent with the enrichment of cytoskeletal remodeling pathways identified by GO analysis. These findings suggest that protracted heroin abstinence is characterized primarily by coordinated suppression of mitochondrial bioenergetic networks, whereas cocaine abstinence is associated with more modest proteomic alterations involving mitochondrial import and cytoskeletal and astrocyte-associated proteins.

### Multiple upstream regulatory pathways converge on bioenergetic dysfunction during heroin abstinence

The broad proteomic remodeling observed during heroin and cocaine abstinence may be driven by alterations in key molecular factors that regulate diverse aspects of cellular biology. To identify potential regulatory mechanisms underlying the proteomic changes observed in mPFC synaptosomes during heroin or cocaine abstinence, we performed IPA upstream regulator analysis of DEPs (**Fig. 2E; Supplementary Table 5**). Consistent with the mitochondrial signature identified by GO enrichment, canonical pathway analysis, and STRING network analysis, upstream regulator analysis predicted coordinated inhibition of regulators governing mitochondrial biogenesis, oxidative metabolism, and bioenergetic substrate availability. Among the highest-ranked predicted inhibited regulators was peroxisome proliferator-activated receptor gamma coactivator-1 alpha (PGC-1α), a master transcriptional coactivator regulating mitochondrial biogenesis and oxidative metabolism ^57, 58^. Frataxin (FXN), a mitochondrial matrix protein required for iron-sulfur cluster assembly and normal activity of electron transport chain Complexes I, II, and III^59^, was similarly predicted to be inhibited. Predicted inhibition of the insulin receptor (INSR) further implicated altered regulation of cellular glucose utilization during heroin abstinence, which was accompanied by protein-level downregulation of glucose transporter 1 (GLUT1). These findings implicate coordinated inhibition of regulatory programs governing mitochondrial biogenesis, respiratory function, and bioenergetic substrate availability during heroin abstinence.

Several activated upstream regulators were associated with mitochondrial quality control and metabolic adaptation. These included the mitochondrial protease CLPP and mitochondrial disaggregase CLPB, which maintain mitochondrial protein integrity^60, 61^, as well as carnitine palmitoyltransferase 1B (CPT1B), which regulates mitochondrial fatty acid import^62^. Although mitochondrial Rho GTPase 2 (MIRO2) was not identified as an upstream regulator, it was the most strongly upregulated mitochondrial protein in the heroin dataset and provides complementary evidence for activation of mitochondrial trafficking and PINK1-Parkin-associated quality-control pathways^63^. RICTOR, a core component of mTOR complex 2^64^, and the histone demethylase KDM5A^65^, were also predicted to be activated, indicating that bioenergetic remodeling was accompanied by broader signaling and epigenetic regulatory programs.

By comparison, cocaine abstinence produced a comparatively muted upstream regulatory profile, with fewer predicted regulators and smaller activation or inhibition magnitudes. Krüppel-like factor 6 (KLF6), a stress-responsive transcription factor^66^, was the strongest predicted activated regulator. In addition, numerous microRNAs were identified as candidate upstream regulators in cocaine-abstinent animals, a pattern that was not prominent following heroin abstinence and that suggests a comparatively greater contribution of post-transcriptional regulation to the cocaine-associated proteomic signature.

Collectively, these findings suggest distinct regulatory landscapes within mPFC synapses during heroin and cocaine abstinence. Heroin abstinence was characterized by predicted inhibition of regulators supporting mitochondrial biogenesis, respiratory function, and glucose utilization, accompanied by predicted activation of mitochondrial quality-control and metabolic-adaptation pathways. In contrast, cocaine abstinence exhibited a more limited regulatory response distinguished by KLF6 activation and enrichment of candidate microRNA regulators. Together with GO enrichment, IPA canonical pathway analysis, and STRING network analysis, the results implicate mitochondrial bioenergetic remodeling as a dominant molecular signature of protracted heroin abstinence and suggest that distinct upstream regulatory mechanisms contribute to the divergent synaptic adaptations across drug classes.

### Whole mPFC metabolomics supports heroin-specific remodeling of bioenergetic pathways

Given the coordinated remodeling of bioenergetic pathways identified by synaptosome proteomics following heroin abstinence, we next asked whether these protein-level alterations were accompanied by corresponding changes in metabolism. A separate cohort of rats underwent long-access IVSA for heroin, cocaine, or saline followed by 30 days of homecage forced abstinence using identical procedures to rats used for proteomics (**Fig. 3A**). Whole mPFC tissue was subsequently collected for targeted LC-MS/MS metabolomic profiling (**Fig. 3A**). Following quality-control filtering, 110 total metabolites were retained for downstream analysis.

**Figure 3.**
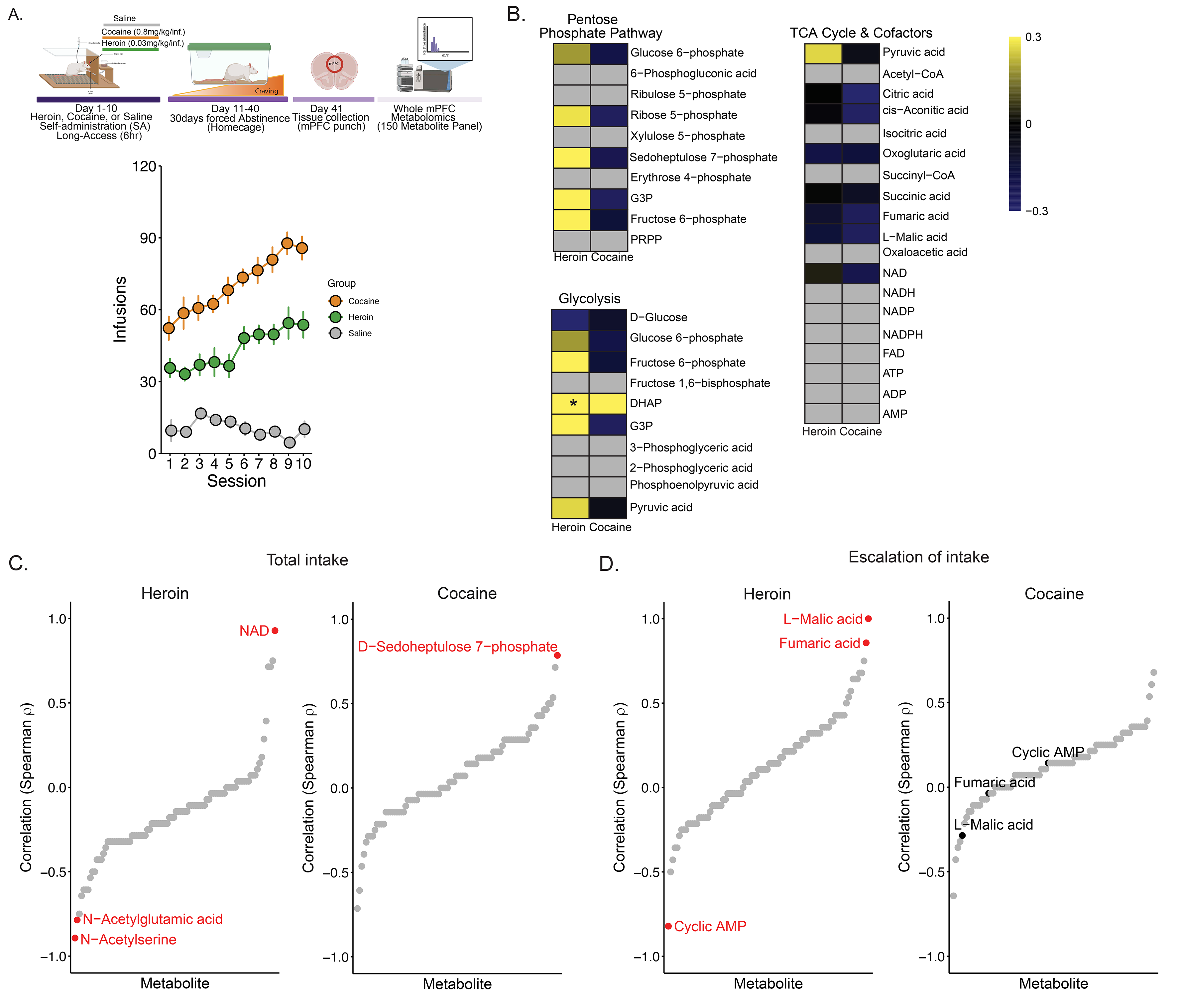
Prolonged heroin abstinence is associated with metabolic remodeling in the mPFC that relates to prior escalation of heroin intake. **A.** Experimental design for targeted metabolomic profiling. Rats underwent 10 days of long-access (6h/day) IVSA of heroin (0.03mg/kg/infusion), cocaine (0.8mg/kg/infusion), or saline, followed by 30 days of home-cage forced abstinence. Whole mPFC tissue was collected on abstinence day 30 for targeted LC-MS/MS metabolomic analysis. Lower panel shows the number of heroin, cocaine, and saline infusions across self-administration sessions, demonstrating higher intake of heroin (Group: F_(1,120)_=386.9, p< 0.001) and cocaine (Group: F_(1,120)_=1179.8, p< 0.001) compared to saline. **B.** Heatmaps showing log_2_ fold changes in metabolite abundance relative to saline controls for metabolites within the pentose phosphate pathway, glycolysis, and tricarboxylic acid (TCA) cycle intermediates and associated redox/energy cofactors. Heroin abstinence was associated with coordinated increases in glycolytic and pentose phosphate pathway intermediates, whereas cocaine abstinence produced comparatively modest metabolic alterations. Dihydroxyacetone phosphate (DHAP) was significantly elevated (log_2_FC = 1.30, *p = 0.019) in heroin abstinence. **C.** Spearman correlation coefficients relating metabolite abundance on abstinence day 30 to total cumulative heroin (left) or cocaine (right) intake during self-administration. Red labels indicate metabolites significantly associated with cumulative drug. In heroin-abstinent animals, NAD was positively associated with cumulative heroin intake (ρ = 0.928, p< 0.01), whereas N-acetylserine and N-acetylglutamic acid were negatively associated (ρ = −0.893, p< 0.01 and ρ = −0.786, p< 0.01, respectively). In cocaine-abstinent animals, D-sedoheptulose 7-phosphate was positively associated with cumulative cocaine intake (ρ = 0.787, p< 0.05). **D.** Spearman correlation analysis relating metabolite abundance on abstinence day 30 to escalation of heroin (left) or cocaine (right) intake during self-administration. Red labels identify metabolites significantly associated with heroin escalation, and the corresponding metabolites are highlighted in the cocaine group for comparison (grey). In heroin-abstinent animals (left), L-malic acid (malate) was found to be significantly positively correlated with escalation of heroin intake (ρ=1.0, p= 0.00002), as was fumaric acid (fumarate; ρ=0.857, p=0.014) while cyclic AMP was negatively associated (ρ=−0.821, p=0.023). No metabolite significantly correlated with escalation of cocaine intake (right).

To determine whether the bioenergetic remodeling identified by synaptosomal proteomics was reflected at the metabolite level, we examined intermediates spanning the three interconnected pathways of central carbon and bioenergetic metabolism: the pentose phosphate pathway (PPP), glycolysis, and the TCA cycle (**Fig. 3B**). Heroin abstinence produced a pattern of directional increases in several PPP and upper glycolytic intermediates that was largely absent following cocaine abstinence. Within the PPP, sedoheptulose 7-phosphate showed increased abundance in heroin-abstinent rats (log_2_FC = 1.16, p = 0.073), accompanied by directional increases in several additional sugar-phosphate intermediates. A similar pattern was observed in glycolysis, where dihydroxyacetone phosphate (DHAP), a downstream intermediate generated following cleavage of fructose-1,6-bisphophate, was significantly elevated (log_2_FC = 1.30, p = 0.019). Although not significant, fructose 6-phosphate (log_2_FC = 0.76, p = 0.105) and glyceraldehyde-3-phosphate (G3P) (log_2_FC =0.42, p=0.121) exhibited similar directional increases. In contrast, metabolites within the TCA cycle remained comparatively stable in both heroin- and cocaine-abstinent animals despite the extensive downregulation of mitochondrial proteins identified by synaptosomal proteomics (**Fig. 2C and D**). These findings provide complementary evidence that prolonged heroin abstinence is associated with remodeling of bioenergetic pathways, characterized by elevated levels of upper glycolytic and PPP intermediates consistent with disruption at the interface between cytosolic carbohydrate metabolism and mitochondrial oxidation.

### Persistent metabolic remodeling is associated with the severity of heroin intake

We next examined whether prolonged metabolic alterations were associated with individual differences in heroin or cocaine exposure. Correlation analyses between cumulative heroin intake with metabolite abundance identified a strong positive relationship between NAD (ρ = 0.928, p< 0.01), a central redox cofactor linking glycolysis, the TCA cycle, and oxidative phosphorylation (**Fig. 3C**, left**; Supplementary Table 6**). Additionally, N-acetylserine and N-acetylglutamic acid each exhibited nominal negative correlations with cumulative heroin intake (ρ = −0.893, p< 0.01 and ρ = −0.786, p< 0.01, respectively). In cocaine-abstinent animals, D-sedoheptulose 7-phosphate showed a significant positive correlation with cumulative cocaine intake (ρ = 0.787, p< 0.05; **Fig. 3C**, right**; Supplementary Table 6**).

One feature of the long-access IVSA procedure is the propensity of animals to exhibit escalating levels of drug intake across sessions, which may reflect changes to underlying neural circuits that promote increasing motivation to consume drug as a hallmark of addiction vulnerability^67–69^. We thus examined whether prior escalation may be associated with a unique pattern of metabolic regulation during protracted abstinence from heroin or cocaine to identify potential metabolic indicators of prior use severity. Rats exhibited progressive increases in drug intake across self-administration sessions 3-10 for both heroin (2.85 inf/session; 95% CI: 1.28-4.42, p = 0.00087) and cocaine (4.08 inf/session; 95% CI: 2.04-6.12, p = 0.00029), confirming escalation of intake (**Supplementary Fig. 2**). We then correlated each individual animal escalation scores with abundance of reliably detected metabolites. In heroin-abstinent rats, L-malic acid (malate) and fumaric acid (fumarate) exhibited a positive correlation with escalation severity (ρ = 0.990, p< 0.001 and ρ = 0.857, p< 0.05, respectively; **Fig. 3D**, left**; Supplementary Table 7**). Malate and fumarate are sequential intermediates in the TCA cycle, and their positive associations with escalation severity identify this metabolic node as potentially related to the degree of prior addiction-like drug intake. Notably, malate dehydrogenase 2 (MDH2), the mitochondrial enzyme catalyzing conversion of malate to oxaloacetate, was significantly reduced in heroin-abstinent synaptosomes (log_2_FC = −0.36, p< 0.001), providing a potential molecular link to altered malate handling. Additionally, cyclic AMP (cAMP) showed a significant negative association with heroin escalation (ρ = −0.821, p< 0.05). In contrast, no metabolite was found to be significantly associated with escalation of cocaine intake (**Fig. 3D**, right**; Supplementary Table 7**). These findings demonstrate that lasting remodeling of bioenergetic pathways within the mPFC is associated with both heroin abstinence and the severity of addiction-relevant patterns of heroin intake.

### Protracted heroin abstinence reduces intrinsic excitability of layer V mPFC pyramidal neurons as well as spontaneous excitatory synaptic transmission

Having identified persistent and heroin-specific proteomic and metabolic remodeling of bioenergetic pathways during heroin abstinence, we next determined whether these molecular changes were accompanied by alterations to mPFC function. Given the relative paucity of bioenergetic remodeling observed in cocaine abstinence, electrophysiological recordings were focused on heroin-abstinent animals. Whole-cell patch clamp electrophysiology recordings were obtained from prelimbic mPFC following 30-day abstinence from heroin or saline IVSA, specifically targeting layer V pyramidal neurons which represent the primary output population of this region (**Fig. 4A and B**). Current-clamp recordings revealed a significant reduction in intrinsic excitability, with layer V pyramidal neurons from heroin-abstinent animals generating fewer action potentials across depolarizing current injections than saline controls (**Fig. 4C**). Heroin abstinence also significantly reduced spontaneous excitatory postsynaptic current (sEPSC) frequency relative to saline controls (**Fig. 4D**), indicating reduced excitatory synaptic drive. These findings show that 30-day heroin abstinence is associated with prolonged reductions in both intrinsic excitability and excitatory synaptic transmission in mPFC layer V pyramidal neurons.

**Figure 4.**
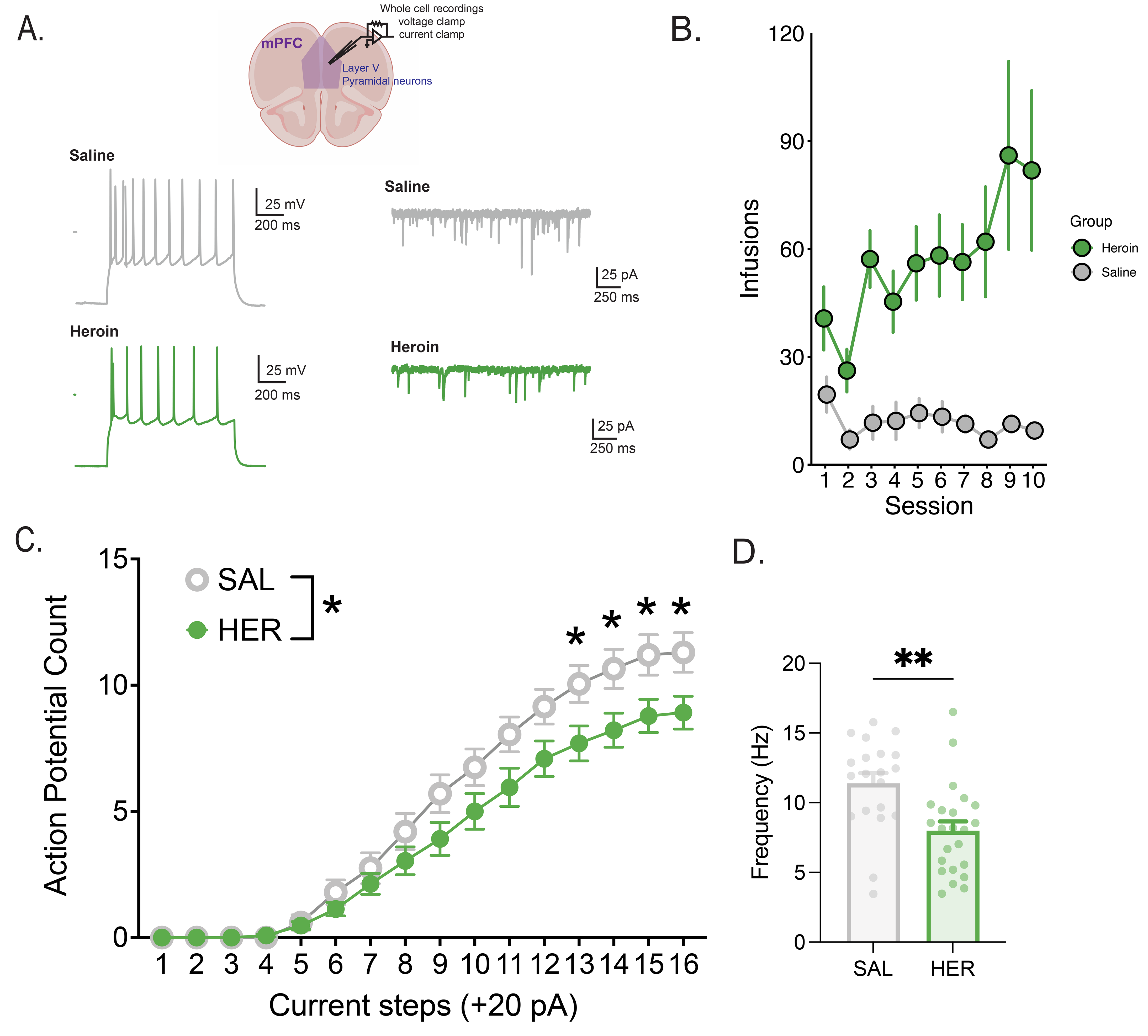
Prolonged heroin abstinence reduces intrinsic excitability and excitatory synaptic drive-in layer V mPFC pyramidal neurons. **A.** Experimental overview and representative whole-cell patch-clamp recordings obtained from layer V pyramidal neurons in the medial prefrontal cortex (mPFC) of rats following 30 days homecage forced abstinence from IVSA of heroin (0.03 mg/kg/infusion) or saline. Representative action potential firing in response to depolarizing current injection recorded under current-clamp conditions (left). Representative spontaneous excitatory postsynaptic currents (sEPSCs) recorded under voltage-clamp conditions (right). Scale bars are indicated in each trace. **B.** Number of heroin or saline infusions earned over IVSA sessions, showing heroin intake was significantly higher than saline intake (Group: F_(1,100)_=96.4, p< 0.001. **C.** Number of action potentials generated in response to increasing depolarizing current injections (+20 pA increments) in layer V mPFC pyramidal neurons from saline (SAL, n=20 neurons from 6 rats) and heroin (HER, n=23 neurons from 6 rats) groups. Heroin abstinence reduced action potential firing across depolarizing current steps, indicating decreased intrinsic neuronal excitability (Two-way ANOVA, F(1,41)=4.595, p=0.038). **D.** Mean spontaneous excitatory postsynaptic current (sEPSC) frequency in layer V mPFC neurons from saline- and heroin-abstinent rats. sEPSC frequency significantly decreased in heroin group compared with saline controls, consistent with reduced excitatory synaptic drive (Welch’s t test, t(40.00)=3.398, p=0.0015). Dots represent individual cells and bars represent mean ± SEM.

### Integrated proteomic and metabolic analysis reveal coordinated remodeling of bioenergetic pathways during heroin abstinence

Having established that 30-day heroin abstinence is associated with coordinated alterations in the synaptic proteome, whole-mPFC metabolism, and neuronal function, we next integrated the proteomic and metabolomic datasets to map these findings into a unified pathway-level framework (**Fig. 5**).Protein meeting the differential abundance criteria and metabolites exceeding the predefined log2 FC threshold were mapped onto major bioenergetic pathways, including the PPP, glycolysis, pyruvate metabolism, TCA cycle, and oxidative phosphorylation. Integrated proteomic and metabolomic analyses reveal more extensive remodeling of energy metabolism following heroin abstinence than cocaine abstinence. In heroin-abstinent animals (**Fig. 5A**), increased abundance of the glycolytic rate-limiting enzymes phosphofructokinase isoforms (PFKL and PFKM), and elevated acetyl-CoA occurred alongside coordinated reductions in pyruvate dehydrogenase complex subunits (PDHA1, PDHB, and DLAT), pyruvate dehydrogenase phosphatase (PDP1), aconitase 2 (ACO2), multiple electron transport chain complex subunits (Complexes I-V), and ATP synthase. These alterations extended from upper glycolysis through pyruvate metabolism and mitochondrial energy-producing pathways, revealing coordinated remodeling of multiple components of cellular bioenergetics. In contrast, cocaine abstinence was associated with comparatively modest molecular alteration (**Fig. 5B**), including increased TALDO1 and DHAP, together with decreased ribose-1-phosphate, L-lactate, PDP1, citrate, cis-aconitate, and Complex I components. Most other proteins and metabolites represented within these bioenergetic pathways showed little change relative to saline controls.

**Figure 5.**
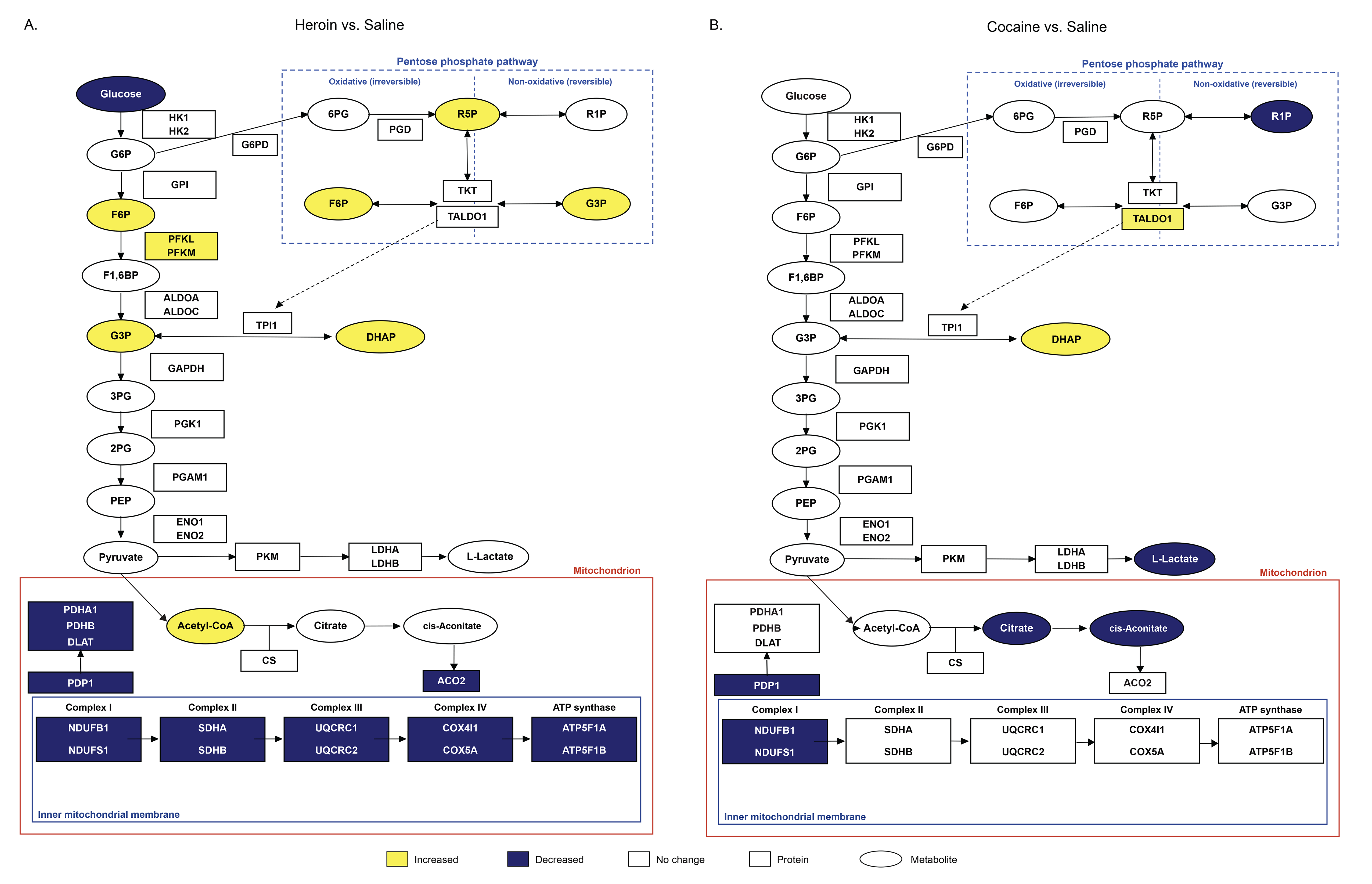
Integrated proteomic and metabolomic analyses reveal more extensive remodeling of energy metabolism in mPFC following heroin than cocaine abstinence. **A-B.** Integrated summary of alterations in central energy metabolism identified by proteomic and whole-mPFC metabolomic analyses following 30 days of abstinence from heroin (**A**) or cocaine (**B**) self-administration. Rectangles represent proteins and ovals represent metabolites. For proteins, yellow and dark blue indicate significantly increased or decreased abundance, respectively (|log fold change| ≥ 0.263, p<0.05), whereas white indicates no significant change relative to saline controls. For metabolites, yellow and dark blue indicate increased or decreased log fold change relative to saline controls, respectively, and white indicates no detectable change. Heroin abstinence was characterized by increased expression of several glycolytic and pentose phosphate pathway intermediates, including fructose 6-phosphate, glyceraldehyde-3-phosphate, dihydroxyacetone phosphate, and ribose 5-phosphate, together with increased PFKL and PFKM protein abundance and increased acetyl-CoA. These changes were accompanied by decreased abundance of pyruvate dehydrogenase complex subunits (PDHA1, PDHB, and DLAT), pyruvate dehydrogenase phosphatase (PDP1), aconitase 2 (ACO2), electron transport chain complexes 1-V, and ATP synthase. Cocaine abstinence produced a more restricted pattern that included increased TALDO1 and dihydroxyacetone phosphate (DHAP), decreased ribose1-phosphate, L-lactate, PDP1, citrate, cis-aconitate, and Complex I, and no significant changes in most other pathway components shown.

Overall, these findings identify coordinated remodeling of bioenergetic pathways as a defining molecular signature of 30-day heroin abstinence in the mPFC, linking prolonged alterations in the synaptic proteome and metabolome with suppression of neuronal excitability. In contrast, cocaine abstinence produced comparatively modest molecular remodeling, indicating that lasting dysfunction of the mPFC during abstinence is associated with fundamentally distinct biological adaptations across these two classes of addictive drugs.

## Discussion

The goal of these studies was to examine biological changes that may underlie mPFC dysfunction during protracted abstinence from drug-taking and contribute to relapse vulnerability. To achieve this, we performed synaptic proteomics and whole-tissue metabolomics in animals undergoing protracted abstinence from IVSA of heroin or cocaine. Heroin abstinence was associated with coordinated remodeling of bioenergetic pathways within the mPFC, characterized by broad reductions in proteins supporting mitochondrial oxidative metabolism together with altered glycolytic and pentose phosphate pathway metabolites. These bioenergetic changes were accompanied by diminished mPFC pyramidal neuron excitability and excitatory synaptic drive measured by patch clamp electrophysiology. Cocaine abstinence, by contrast, produced a more limited molecular footprint that shared some features with heroin abstinence but was substantially less extensive and less coordinated. Taken together, these findings identify persistent bioenergetic remodeling as a prominent molecular signature of opioid abstinence that was substantially less evident following cocaine abstinence, providing a candidate substrate that may contribute to sustained mPFC dysfunction associated with relapse vulnerability.

### A convergent bioenergetic phenotype in mPFC during protracted abstinence

Mitochondria are concentrated at synapses to meet the substantial ATP demands of neurotransmission, and disruption of mitochondrial homeostasis can impair sustained synaptic function with impacts on cognition and behavior^33, 70, 71^. Here, we find that the dominant molecular feature of the mPFC synaptic proteome following protracted heroin abstinence was a coordinated and comprehensive remodeling of mitochondrial bioenergetic pathways (**Fig. 2**). These changes were widespread, with reductions in proteins spanning multiple stages of mitochondrial ATP production, including the PDH complex, TCA cycle enzymes, electron transport chain ETC complexes I-IV, ATP synthase, mitochondrial protein import machinery, and mitochondrial chaperones. Upstream regulator analysis predicted PGC-1α, a master coactivator of mitochondrial biogenesis and oxidative metabolism^58^, to be inhibited, which may provide a regulatory mechanism through which this coordinated reduction in mitochondrial proteins could arise. Notably, this remodeling was not uniformly suppressive: MIRO2, a mitochondrial outer-membrane GTPase involved in mitochondrial trafficking and quality control^72^, was the most upregulated protein detected, suggesting that mitochondrial maintenance pathways may be engaged in parallel with remodeling of bioenergetic machinery.

Our whole-mPFC metabolomics analysis provided complementary evidence for bioenergetic remodeling during heroin abstinence, although these changes were more selective (**Fig. 3; Supplementary Tables 6 and 7**). We found an accumulation of upstream glycolytic and PPP intermediates, with significantly elevated DHAP, and directional increases in fructose-6-phosphate, glyceraldehyde-3-phosphate, and sedoheptulose-7-phosphate. Because metabolomics measures steady-state metabolite abundance rather than pathway flux^73^, these changes cannot distinguish increased glycolytic activity from reduced downstream utilization. However, a potential point of convergence with the synaptic proteomics results is the coordinated reduction of all three subunits of the PDH complex (PDHA1, PDHB, and DLAT), which controls carbon entry from glycolysis into mitochondrial oxidation by converting pyruvate to acetyl-CoA^74^ (**Fig. 5A**). Reduced PDH complex activity would be expected to produce the upstream accumulation of glycolytic intermediates observed here, providing a mechanistic bridge between the two datasets. It is worth noting that this reduction in PDH complex subunits was accompanied by increased, rather than decreased, whole tissue acetyl-CoA abundance (**Fig. 5A**). Compensatory acetyl-CoA production via fatty-acid β-oxidation, consistent with the predicted activation of CPT1B, might cause this discrepancy but further study is needed. More broadly, because the proteomic and metabolomic datasets were generated from different biological compartments and cohorts, using synaptosome-enriched preparations versus whole-tissue homogenates containing non-synaptic mitochondria and glia, the apparent convergence between the two datasets should be regarded as suggestive rather than confirmatory and does not establish that the same molecular pools are being altered. However, TCA intermediates were relatively unchanged in heroin or cocaine abstinence, and several bioenergetic cofactors were below reliable detection thresholds, likely reflecting dilution of synapse-specific changes in this whole-tissue analysis. These findings emphasize the utility of integrating compartment-enriched proteomic analyses with whole-tissue metabolomics while also highlighting the need for direct measurements of metabolic flux and mitochondrial function.

In parallel with energetic changes, both proteomics and metabolomics datasets converged on pathways supporting cellular redox homeostasis (**Fig. 5**). Glutathione metabolism was the most enriched biological process among proteins increased in heroin-abstinent synaptosomes, including GCLC, the rate-limiting enzyme of glutathione biosynthesis^75^. In addition, several PPP intermediates showed directional increases in the metabolome; the oxidative PPP is a major source of NADPH required for regeneration of reduced glutathione^76^. Reduced abundance of mitochondrial Complex I proteins may also influence NAD+ dependent redox metabolism, as Complex I is the primary site of NADH oxidation within the electron transport chain and contributes to cellular NAD+/NADH balance^77^. Complementing this, NAD+ abundance was associated with the magnitude of heroin intake in the metabolome, while IPA identified enrichment of sirtuin signaling – an NAD+ dependent pathway that couples cellular energetic and redox status to transcriptional regulation and stress responses^78^ and has been implicated previously in other brain regions in opioid and cocaine action^79^. These findings suggest that 30-day heroin abstinence is accompanied by remodeling across multiple interdependent arms of redox metabolism, spanning glutathione-dependent antioxidant defense and NAD-dependent signaling. These bioenergetic and redox adaptations may contribute to the persistent mPFC hypofunction widely reported during prolonged opioid abstinence (22011681) by limiting the energetic capacity required to sustain synaptic transmission and neuronal firing. Consistent with this possibility, heroin abstinence in the present study was accompanied by reduced intrinsic excitability and decreased excitatory synaptic drive in mPFC pyramidal neurons (**Fig. 4**), providing a functional correlate of the molecular remodeling identified here. These findings build on prior work demonstrating that repeated opioid exposure alters glutathione-dependent redox homeostasis through reductions in glutathione levels and glutathione-dependent enzyme activity in both experimental models and human tissue^80, 81^. Recent proteomic studies of postmortem human brains from individuals with OUD also point to altered oxidative stress-associated molecular signatures^82^. In contrast to prior studies, our findings of increased glutathione biosynthetic machinery one month into abstinence raise the possibility that sustained or compensatory engagement of antioxidant pathways persists beyond active drug exposure. However, we did not reliably quantify GSH/GSSG, NADH, or NADPH, and future work will be required to establish whether oxidative stress, antioxidant capacity, or cellular redox state is net increased or decreased during prolonged heroin abstinence.

### Bioenergetic remodeling in mPFC may scale with escalation of drug intake

Escalation of drug intake observed using extended-access IVSA sessions is thought to model a progressive loss of control over consumption, a behavioral trajectory considered to be central to the development of addiction-like behavior^67, 83, 84^. We therefore examined whether the severity of intake escalation of heroin or cocaine was associated with distinct metabolic signatures in the mPFC during protracted abstinence. This analysis revealed a strong positive correlation between escalation of heroin intake and the abundance of malate (**Fig. 3C**), suggesting that metabolic remodeling of the TCA cycle scales with the severity of prior heroin intake. Fumarate, the immediate precursor of malate in the TCA cycle, was also positively associated with heroin escalation at the nominal level. Cyclic AMP was negatively associated with heroin escalation at the nominal level. In contrast, no metabolites were significantly associated with escalation of cocaine intake.

Interestingly, our synaptic proteomic analysis identified reduced abundance of MDH2, the enzyme catalyzing malate’s conversion to oxaloacetate, providing a potential mechanistic link to altered malate handling^85^ (**Fig. 5A**). However, malate did not differ significantly between heroin and saline groups, and the proteomic and metabolomic measurements were obtained from different biological compartments; therefore, these findings do not establish substrate accumulation caused by an MDH2 bottleneck. Instead, the convergence across independent datasets identifies this node of the TCA cycle as a candidate metabolic adaptation associated with the severity of prior heroin intake escalation.

### Bioenergetic remodeling in mPFC during heroin abstinence is accompanied by reduced mPFC neuronal excitability

Synaptic transmission is extremely energetically demanding, necessitating continuous ATP availability to sustain ion gradients, neurotransmitter release, and vesicle recycling^33, 86^. Thus, it is plausible that the coordinated remodeling of mitochondrial bioenergetic pathways in mPFC would be accompanied by altered neuronal function during prolonged heroin abstinence. Consistent with this prediction, mPFC layer V pyramidal neurons in heroin-abstinent rats generated fewer action potentials in response to depolarizing current injection and displayed reduced sEPSC frequency, indicating reduced intrinsic membrane excitability and excitatory synaptic drive (**Fig. 4D**). Because layer V pyramidal neurons constitute a major output population of the mPFC^87^, these findings provide functional evidence that 30-day heroin abstinence is associated with lasting suppression of pyramidal neuron excitability. Prior work has established that AMPA receptor plasticity at mPFC synapses is recruited during cue-induced heroin relapses, with GluR2 internalization producing synaptic depression that is mechanistically necessary for reinstatement of heroin seeking^88^. More recently, reduced AMPAR- and NMDAR-mediated synaptic transmission in prelimbic pyramidal neurons has been observed at 14 days of heroin abstinence alongside suppressed basal excitatory neuron activity in vivo, demonstrating that mPFC synaptic depression emerges early in the abstinence period^89^. Our findings extend this work by demonstrating that deficits in intrinsic excitability and excitatory synaptic drive persist through one month of abstinence, suggesting that reduced cortical excitability may represent a sustained neurophysiological state. Further, our results demonstrate that mPFC pyramidal neurons are suppressed at baseline during protracted abstinence, in the absence of any drug-associated stimuli, suggesting that opioid abstinence leaves the mPFC in a persistently hypoexcitable state that may compromise its capacity to suppress drug-seeking in response to relapse-triggering stimuli. Although we did not directly test whether bioenergetic remodeling drives reduced neuronal excitability, the coordinated reductions in proteins supporting oxidative metabolism and accompanying metabolic remodeling observed in independent cohorts undergoing identical procedures provide a plausible biological context for the lasting neuronal hypofunction observed here.

### Heroin and cocaine engage distinct molecular mechanisms during protracted abstinence

Despite their shared ability to induce persistent relapse-relevant mPFC dysfunction^15^, we observed substantially more extensive molecular remodeling following protracted abstinence from heroin compared to cocaine (**Fig. 1D-F**). Within the mPFC synaptic proteome, cocaine abstinence was characterized by reductions in mitochondrial import and processing machinery, including TIMM44, TOMM22, and mitochondrial-processing peptidase subunit alpha (PMPCA)^52^, directionally consistent with heroin but considerably narrower in scope (**Fig. 2D**). The upregulated signature pointed to structural reorganization, including axonal cytoskeletal proteins NFL and NFM^90^, together with ELAVL4, an RNA-binding protein that stabilizes neuronal transcripts involved in axonal growth and cytoskeletal remodeling^53^, raising the possibility that similar post-transcriptional mechanisms could contribute to the concurrent increase we observed in NFL and NFM, and the presynaptic active zone scaffolding protein PPFIA3^91^. These data suggest that cocaine abstinence may be biased towards structural remodeling involving axonal and presynaptic components rather than the broad bioenergetic remodeling observed following heroin abstinence.

Notably, downregulation of a shared astrocyte-associated protein network including AQP4, Kir4.1, and connexin-43^92^ was observed in both heroin- and cocaine-abstinent animals, representing a point of convergence between otherwise divergent proteomic signatures, though with cocaine this network was connected with upregulated cytoskeletal proteins. However, STRING connectivity alone cannot establish functional coupling between these processes. Unlike heroin, where mitochondrial bioenergetic and redox-associated changes formed a more coherent molecular signature, no equivalent pathway-level organization was evident following cocaine abstinence. This comparatively limited molecular footprint raises the possibility that persistent cocaine-induced adaptations occur through molecular processes, cell populations, or circuits not fully captured by synaptosomal proteomics or whole-tissue metabolomics. Alternatively, together with previous evidence that both heroin and cocaine produce persistent mPFC dysfunction^15^, our findings suggest that these drug classes may converge on similar circuit-level phenotype through fundamentally distinct molecular mechanisms, with heroin preferentially disrupting mitochondrial bioenergetics and cocaine favoring structural and synaptic remodeling. If so, comparable circuit-level dysfunction may arise through drug class-specific molecular mechanisms, with important implications for developing targeted therapies for relapse vulnerability. These findings support the that opioid and psychostimulant abstinence engage distinct neurobiological programs despite converging on persistent dysfunction of the mPFC.

## Conclusions

The present studies identify previously underappreciated bioenergetic remodeling during protracted heroin abstinence that occurs alongside sustained reductions in mPFC neuronal excitability and may contribute to impaired cortical function associated with relapse vulnerability. Using proteomics and metabolomics across cellular compartments, these findings converge on robust and coordinated mitochondrial remodeling in mPFC synapses during heroin abstinence, accompanied by broader shifts in cytosolic carbohydrate metabolism across the mPFC and engagement of glutathione-mediated antioxidant processes consistent with compensatory responses to cellular stress. These molecular adaptations were accompanied by reduced mPFC pyramidal neuron excitability and excitatory synaptic drive, providing a potential functional link between persistent molecular remodeling and cortical hypofunction during abstinence. These adaptations were largely absent following cocaine abstinence, indicating divergence on the biological basis of relapse vulnerability across opioid and psychostimulant use disorders. These findings suggest mitochondrial bioenergetics and redox homeostasis as candidate biological pathways for future studies aimed at determining whether restoration of metabolic function can reduce lasting cortical dysfunction and relapse vulnerability in opioid use disorder.

## Acknowledgments

We thank NYU Langone Health’s Metabolomics Laboratory for its help in acquiring and analyzing the data presented, and Dr. TuKiet Lam and the Yale/NIDA Neuroproteomics Center (P30 DA018343) for processing the proteomics samples, providing technical support, funding opportunities, and creating proteomics repository. Illustration of experimental design was created with BioRender.com.

---

**Supplementary Figure 1. Behavioral validation and global proteomic changes following heroin and cocaine self-administration. A-B.** Active and inactive lever presses across heroin (A), cocaine (B), and saline self-administration sessions (n=7 rats/group). Heroin-trained and cocaine-trained rats developed preferential responding on the active lever. **C.** Principal component analysis (PCA) of mPFC synaptosomal proteomic profiles from saline-, heroin-, and cocaine-exposed rats after 30 days of withdrawal. Principal components (PC) 1 and 2 accounted for 19.2% and 14.0% of the total variance, respectively, and illustrate the distribution of samples according to treatment conditions. **D.** Heatmap of z-score–normalized mean protein abundance for saline, heroin, and cocaine groups. Each row represents a protein, and each column represents the mean abundance of a treatment group. Red indicates higher relative abundance, and blue indicates lower relative abundance across group means.

**Supplementary Figure 2. Escalation of heroin and cocaine intake across long-access self-administration sessions.** Total drug infusions earned across self-administration sessions (S3– S10) in rats trained to self-administer heroin (2.85 inf/session; 95% CI: 1.28–4.42, p= 0.00087, left) or cocaine (4.08 inf/session; 95% CI: 2.04–6.12, p = 0.00029, right). Gray lines represent individual animals, and colored symbols represent group means ± SEM. Both heroin and cocaine groups exhibited significant progressive increases in drug intake across sessions 3-10, confirming escalation under long-access self-administration conditions.

**Supplementary Table 1. Differentially expressed proteins in mPFC synaptosomes following cocaine or heroin abstinence**. Filtered lists of differentially expressed proteins (DEPs) identified in mPFC synaptosomes following 30 days of abstinence from cocaine or heroin self-administration relative to saline controls. Separate worksheets contain the Heroin vs. Saline (HvS) and Cocaine vs. Saline (CvS) comparisons. DEPs were defined using a nominal limma p< 0.05 and a ≥20% change in protein abundance (|log FC|>0.263). Columns include UniProt accession (UniProt_Code), fold change (FC), average log protein abundance (AveExpr), moderated *t* statistic (*t*), nominal limma *P* value (pvalue), Benjamini–Hochberg-adjusted p value (adj.P.Val), log-odds of differential expression (B), empirical *P* value (empirical_pval), protein annotation (ProteinInfo and Protein_Name), and gene symbol (GeneName). Empirical p values are provided as supplementary statistical information but were not used to define differential protein abundance.

**Supplementary Table 2. Gene Ontology enrichment analysis of proteins altered following heroin abstinence**. Complete GO enrichment results for proteins differentially expressed in mPFC synaptosomes following 30 days of abstinence from heroin self-administration relative to saline controls. Separate worksheets contain results from the 2025 GO Biological Process (GO_BP), Cellular Component (GO_CC), and Molecular Function (GO_MF) databases. Enrichment analyses were performed separately for increased and decreased proteins, as indicated by the Direction column. Columns include GO term, overlap, nominal P value, adjusted P value, Enrichr legacy P values, odds ratio, combined score, contributing genes, comparison, direction of protein abundance change, GO category, and −log10(p value).

**Supplementary Table 3. Gene Ontology enrichment analysis of proteins altered following cocaine abstinence**. Complete GO enrichment results for proteins differentially expressed in mPFC synaptosomes following 30 days of abstinence from cocaine self-administration relative to saline controls. Separate worksheets contain results from the 2025 GO Biological Process (GO_BP), Cellular Component (GO_CC), and Molecular Function (GO_MF) databases. Enrichment analyses were performed separately for increased and decreased proteins, as indicated by the Direction column. Columns include GO term, overlap, nominal P value, adjusted P value, Enrichr legacy P values, odds ratio, combined score, contributing genes, comparison, direction of protein abundance change, GO category, and −log10(p value).

**Supplementary Table 4. Filtered IPA canonical pathway analysis following heroin or cocaine abstinence**. Filtered IPA canonical pathways identified from differentially expressed proteins in mPFC synaptosomes following 30 days of abstinence from heroin or cocaine self-administration relative to saline controls. Separate worksheets contain the Heroin vs. Saline and Cocaine vs. Saline comparisons. Columns include Ingenuity Canonical Pathway, −log10(p value), pathway ratio, activation z score, and molecules contributing to pathway enrichment.

**Supplementary Table 5. Filtered IPA upstream regulator analysis following heroin or cocaine abstinence**. Filtered IPA upstream regulator predictions based on differentially expressed proteins in mPFC synaptosomes following 30 days of abstinence from heroin or cocaine self-administration relative to saline controls. Separate worksheets contain the Heroin vs. Saline and Cocaine vs. Saline comparisons. Columns include upstream regulator, expression log ratio, molecule type, predicted activation state, activation z-score, p value of overlap, target molecules represented in the dataset, and mechanistic network information.

**Supplementary Table 6. Correlations between metabolite abundance and cumulative drug intake**. Correlation analysis relating to mPFC metabolite abundance after 30 days of abstinence to cumulative heroin or cocaine intake during self-administration. Separate rows report results for each metabolite within the heroin and cocaine groups. Columns include metabolite variable, Pearson correlation coefficient (r□) and corresponding p value, Spearman correlation coefficient (r□) and corresponding p value, effective sample size (neff), drug group, and Benjamini– Hochberg FDR-adjusted q value for the Spearman correlation (q□).

**Supplementary Table 7. Correlations between metabolite abundance and escalation of drug intake.** Correlation analysis relating to mPFC metabolite abundance after 30 days of abstinence to individual differences in escalation of heroin or cocaine intake during self-administration. Separate rows report results for each metabolite within the heroin and cocaine groups. Columns include metabolite variable, Pearson correlation coefficient (r□) and corresponding p value, Spearman correlation coefficient (r□) and corresponding p value, effective sample size (neff), drug group, and Benjamini–Hochberg FDR-adjusted q value for the Spearman correlation (q□).

